# Global Neuron Shape Reasoning with Point Affinity Transformers

**DOI:** 10.1101/2024.11.24.625067

**Authors:** J. Troidl, J. Knittel, W. Li, F. Zhan, H. Pfister, S. Turaga

## Abstract

Connectomics is a field of neuroscience that maps the brain’s intricate wiring diagram. Accurate neuron segmentation from microscopy volumes is essential for automating connectome reconstruction. However, state-of-the-art algorithms use image-based convolutional neural networks limited to local neuron shape context. Thus, we introduce a new framework that reasons over global neuron shape with a novel point affinity transformer. Our framework embeds a (multi-)neuron point cloud into a fixed-length feature set from which we can decode any point pair affinities, enabling clustering neuron point clouds for automatic proofreading. We also show that the learned feature set can easily be mapped to a contrastive embedding space that enables neuron type classification using a simple classifier. Our approach excels in two demanding connectomics tasks: correcting segmentation errors and classifying neuron types. Evaluated on three benchmark datasets derived from state-of-the-art connectomes, our method outperforms point transformers, graph neural networks, and unsupervised clustering baselines.

## 1 Introduction

The field of connectomics maps the wiring diagrams of biological neural networks using high-resolution 3D microscopy and image segmentation. A landmark achievement was the full fruit fly brain connectome [8], which required extensive manual proofreading despite advanced segmentation algorithms: 622 researchers contributed 33 human-years to correct 10^5^ neurons in the millimeter-scale Drosophila brain. Scaling up connectomics to map the 500*×* larger mouse brain [1] will require orders of magnitude larger human effort with current technology. Thus, improving automated segmentation accuracy is crucial for advancing neuroscience. Existing segmentation methods [17, 13, 25, 31, 29, 30, 42, 46] are local and CNN-based, lacking the ability to reason over full and sparse neuron shapes. While these methods achieve high accuracy, with error-free paths up to 1.1 mm [17], manual proofreading remains essential for large datasets, such as the 145 meters of wiring in a fruit fly brain [8]. Human proofreaders rely on 3D neuron renderings, which span massive volumes (up to 10^12^ voxels), beyond the practical scope of CNNs. Hence, we propose a point affinity transformer that reasons over global neuron shape using point clouds.

Point clouds efficiently represent sparse, expansive neuron shapes. We frame neuron shape learning as a pairwise affinity prediction task: two points have affinity 1 if they belong to the same neuron, and 0 otherwise. Our architecture avoids explicit spatial features (e.g., grid-based), enabling flexible representation of sparse neuron structures. We apply our method to automated neuron error correction—detecting and fixing point cloud errors—and to neuron type classification via contrastive embeddings. Both tasks require global morphology understanding. We evaluate on three connectome datasets [8, 44, 38], comparing against GNNs [48, 43], other transformers [57, 50], and unsupervised clustering baselines. Our model outperforms these baselines in both tasks across standard metrics.

In summary, our contributions are threefold: *(1)* We introduce a novel neuron shape learning framework based on pairwise point affinity prediction. Given a (multi-)neuron point cloud, our Preprint. Under review. transformer-based model, the point affinity transformer, predicts affinities between point pairs to separate neurons from background (Fig. 2b,c). *(2)* We apply this model using predicted affinities and agglomerative clustering to correct simulated reconstruction errors for which ground truth data exists. Our method outperforms GNNs and other point cloud transformers, as well as unsupervised clustering. *(3)* The model’s internal representations, trained solely on affinity prediction, can be projected into a contrastive embedding space for neuron type classification.

**Figure 1:**
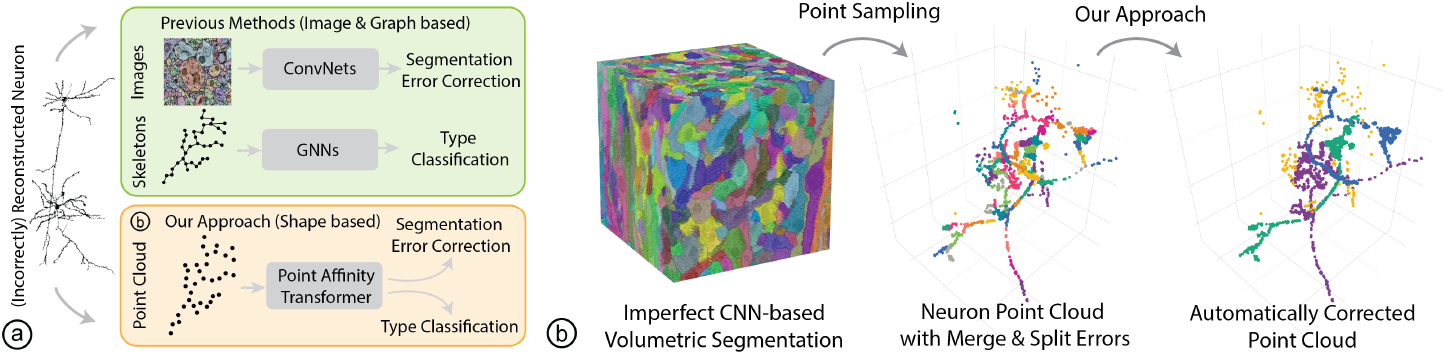
Overview & Workflow. ⓐPrevious approaches use 3D ConvNets for improving neuron segmentation accuracy and are thus limited to local fields of view. ⓑ We propose a framework for neuron segmentation error correction and type classification. 3D CNN-based volumetric neuron segmentations contain errors. We derive a neuron point cloud containing split and merge errors from the connectome segmentation volume. Our approach enables the automatic correction of errors in such erroneous neuron point clouds.

## 2 Related Work

### Point Clouds

We choose point clouds over other spatial representations because they suit sparse, thin structures better than volumetric data and are easily extracted from segmentations without extra processing [21]. Point cloud processing methods include GNNs [48, 43], transformers [57, 51, 50, 56, 55, 14], and set-based models [37, 36]. Transformers are better suited for capturing global neuron shapes, as GNNs struggle with long-range feature updates in sparse data, while self-attention enables distant point interactions. Existing point cloud transformers have limitations. Some [54, 57] rely on spatial references in internal features, limiting flexibility for sparse data. Our model uses a spatially agnostic internal space. Others [57] require fixed input sizes, while our masked attention supports variable-size point clouds. Finally, instance segmentation models like PTv3 + PG [50, 19] assume compact objects, unsuitable for thin, intertwined neurons where centroids are uninformative.

### Neuron Error Correction

Substantial ongoing efforts are manually correcting segmentation errors in large-scale connectomes [8, 9, 10, 41, 32]. Another approach is to reduce the number of errors by increasing the robustness and accuracy of image-based segmentation models [31, 29, 17, 13, 16, 58, 15]. However, these 3D convolutional networks lack the global context of neuron morphology due to inherently localized convolution operations. Instead, we show that point clouds are a robust and expressive 3D representation that allows the global neuron shape context to be considered. Prior work has made progress toward automatically proofreading connectome reconstructions. However, these approaches either rely on handcrafted heuristics [20], build on brittle auxiliary graphs [2, 33, 5], or use constrained local fields of view [39, 58, 16].

### Neuron Classification

Neuron classification methods fall into two categories. The first targets sub-compartment classification [7, 27], labeling structures like dendrites, axons, or somas without needing global context. The second, neuron typing [3, 40, 18, 4, 24], assigns a functional label to the entire neuron, requiring global morphology. Traditional approaches rely on handcrafted features [4, 23], while learning-based methods use skeletons [3, 12, 52], connectome graphs [28], or 2D projections [40], which may not preserve 3D shape.

**Figure 2:**
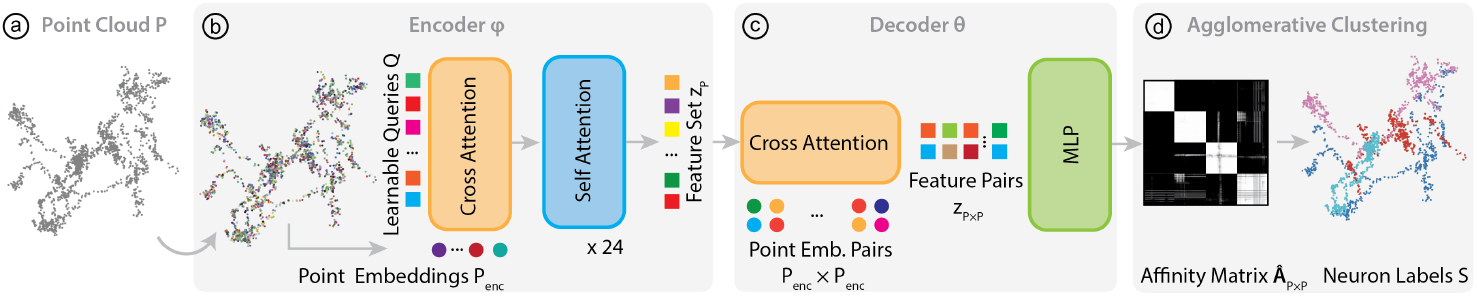
Global Shape Reasoning Pipeline. ⓐOur model encodes a point cloud with samples from *N* neurons ⓑ into a fixed-length feature set using a series of cross- and self-attention layers (Sec. 3.2). ⓒ Next, we condition the decoder with this feature set and compute feature pairs of points from the input point cloud using a cross-attention module. These feature pairs are then decoded into affinity values using an MLP (Sec. 3.2). ⓓ We then extract neuron labels by performing feature agglomerative clustering on the affinity matrix (Sec. 3.3).

## 3 Method

Our goal is to learn an expressive feature space that can capture global neuron shapes. We demonstrate this capability in two challenging connectomics tasks: automated reconstruction error correction using our point affinity transformer model and neuron type classification.

### 3.1 Automated Error Correction Workflow

3D CNNs for neuron segmentation generate voxel-based label volumes, where each label typically represents a “supervoxel”, a small sub-segment of a neuron. Therefore, reconstructing the complete structure of a neuron requires agglomerating numerous supervoxels (Fig.1 ⓐ). The specifics of the segmentation technique determine the approach to agglomeration; however, this process is generally susceptible to segmentation errors, resulting in either split or merge errors. The goal of proofreading is to correct these errors by separating incorrectly merged segments and joining incorrectly split ones. In practice, the balance between split and merge errors is influenced by the level of supervoxel agglomeration. We assume that the model input contains points from multiple neurons. The point cloud may entirely cover some neurons, while others are only partially covered, so-called background fragments (Fig. 1 ⓑ). We sample points from skeleton vertices and discard edge information. Our algorithm then focuses on recoloring the point cloud into individual neurons (Fig. 1 ⓒ) while excluding any background fragments that do not correspond to full neurons.

### 3.2 Point Affinity Transformer

#### Problem Formulation

The main idea of our approach is to formulate global neuron shape learning as a pair-wise affinity prediction task on multi-neuron point clouds. Given an input point cloud *P* = {*p*_0_, …, *p*_*n*_} ⊆ R3 (Fig.2 ⓐ) sampled from {1, …, *K*} neurons and a set of background fragments, our model *M* learns to predict an affinity matrix **Â** *P ×P* ∈ ℝ*n×n* between all point pairs *P × P* as below:

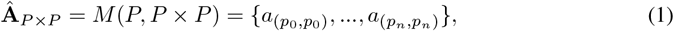

where an affinity value *a*_(*pi,pj*)_ represents the likelihood that *p*_*i*_ and *p*_*j*_ belong to the same neuron. Given **Â** *P* _*×*_ *P*, we reconstruct segmentation labels *S* = {*s*_0_, …, *s*_*n*_}, *s*_*i*_ ⊆ N. *s*_*i*_ ∈ *S* maps to the neuron identity of the respective point *p*_*i*_ ∈ *P*. Specifically, *s* = 0 denotes a background class, which covers fragments that do not assemble entire neurons. The goal is to learn a mapping such that all points with the same semantic label *s >* 0 belong to the same neuron. We derive *S* by clustering *P* using an affinity-based distance metric 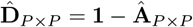 (Sec. 3.3).

For model training, we use the binary cross entropy (BCE) loss between the predicted matrix **Â** and the binary ground truth matrix **A** ∈ {0, 1}^*n×n*^, which we derived from proofread connectome data:

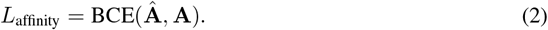

We reduce computational cost during training by decoding a randomly selected subset of point pairs (*P × P*)^*′*^ ⊂ (*P × P*) and thus also only predict a subset of the entire affinity matrix **Â** _(*P × P*)*′*_ ⊂ **Â** _*P × P*_. At test time, we decode point pairs *P × P* to obtain the segmentation label set *S* for points *P*. Alternatively, we can also condition *M* using a query point *q* ∈ *P*, allowing us to compute affinities for a single neuron or background fragment **Â** *P* _*× q*_ = *M* (*P, q*). We discuss the scalability in the supplement (Sec. E).

#### Point Cloud Encoding

Inspired by recent progress in neural fields [55], our point cloud encoding procedure consists of three steps. First, we project each point *p* using a positional frequency encoding *γ* [45] and a fully connected layer (FC) into *D*-dimensional embedding space as below:

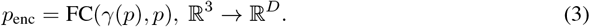

We denote the encoded point cloud as *P*_enc_. Second, *P*_enc_ is transformed into feature set representation *z*^(0)^ using cross-attention [47] with a learnable queries set *Q* ∈ R*D×C* [55] and *P*_enc_ ∈ R*D×*|*P* | as the key-value pair:

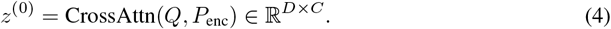

Cross-attention is defined as:

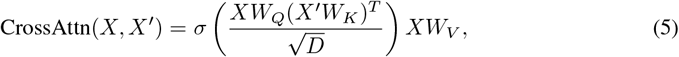

with *X* and *X*^*′*^ being two input sequences, *W*_*Q*_, *W*_*K*_, and *W*_*V*_ denoting three trainable weight matrices, and *σ* representing the *softmax* function. Here, *D* is the dimensionality of all *C* features in the feature set. *Q* is a randomly initialized trainable parameter in our model. Third, we enable information exchange between all elements *z*_*i*_ in the feature set through a series of *j* = 24 sequential self-attention modules:

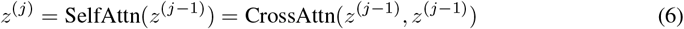

Here, *z*^(*j*)^ is a feature representation of a (multi-)neuron point cloud after the *j* self-attention layer. From now on, we use *z*^(24)^ = *z*_*P*_ for clarity. Note that, unlike other methods [35, 54], *z*_*P*_ has no explicit spatial reference and is thus well suited to fit thin and sparse spatial structures such as neurons flexibly. In summary, the encoder *φ* uses a series of cross- and self-attention layers to encode *P* into a feature set *z*_*P*_ (Fig. 2ⓑ):

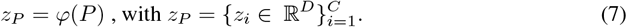

In the following sections, we show how to decode *z*_*P*_ into point pair affinities, which are subsequently used for guiding clustering that leads to the label set *S*.

#### Affinity Decoding

Next, we input the feature set *z*_*P*_ and point embedding pairs *P*_enc_ *× P*_enc_ into the decoder *θ* to predict the affinity matrix **Â** *P ×P* (Fig 2 ⓒ):

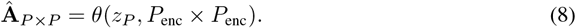

We can either use all point pairs *P × P* to receive the full affinity matrix **Â** *P ×P*, or only use a subset of point pairs (*P × P*)^*′*^ ⊂ (*P × P*). Next, a cross-attention module maps the feature set *z*_*P*_ ∈ R*D×C* into a set of feature pairs *z*_*P ×P*_ ∈ R2*D×*|*P ×P* | given *P*_enc_ *× P*_enc_:

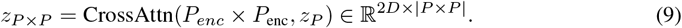

Here, |*P × P* | represents the number of point pairs. Finally, feature pairs *z*_*P ×P*_ are decoded into the affinity matrix **Â** *P* _*×*_ *P* using a MLP : R2*D*→ R. The MLP has a sigmoid final layer to ensure all affinity values are in [0, 1].

### 3.3 Affinity-guided Point Clustering

We derive the set of segmentation labels *S* from the predicted affinity matrix **Â** *P × P*. Since affinities are computed per point pair, we interpret affinity values as a point proximity metric. Thus, we can cluster the point cloud *P* according to an affinity-based distance metric 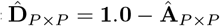 *P ×P* = **1.0** − **Â** *P ×P*.

**Figure 3:**
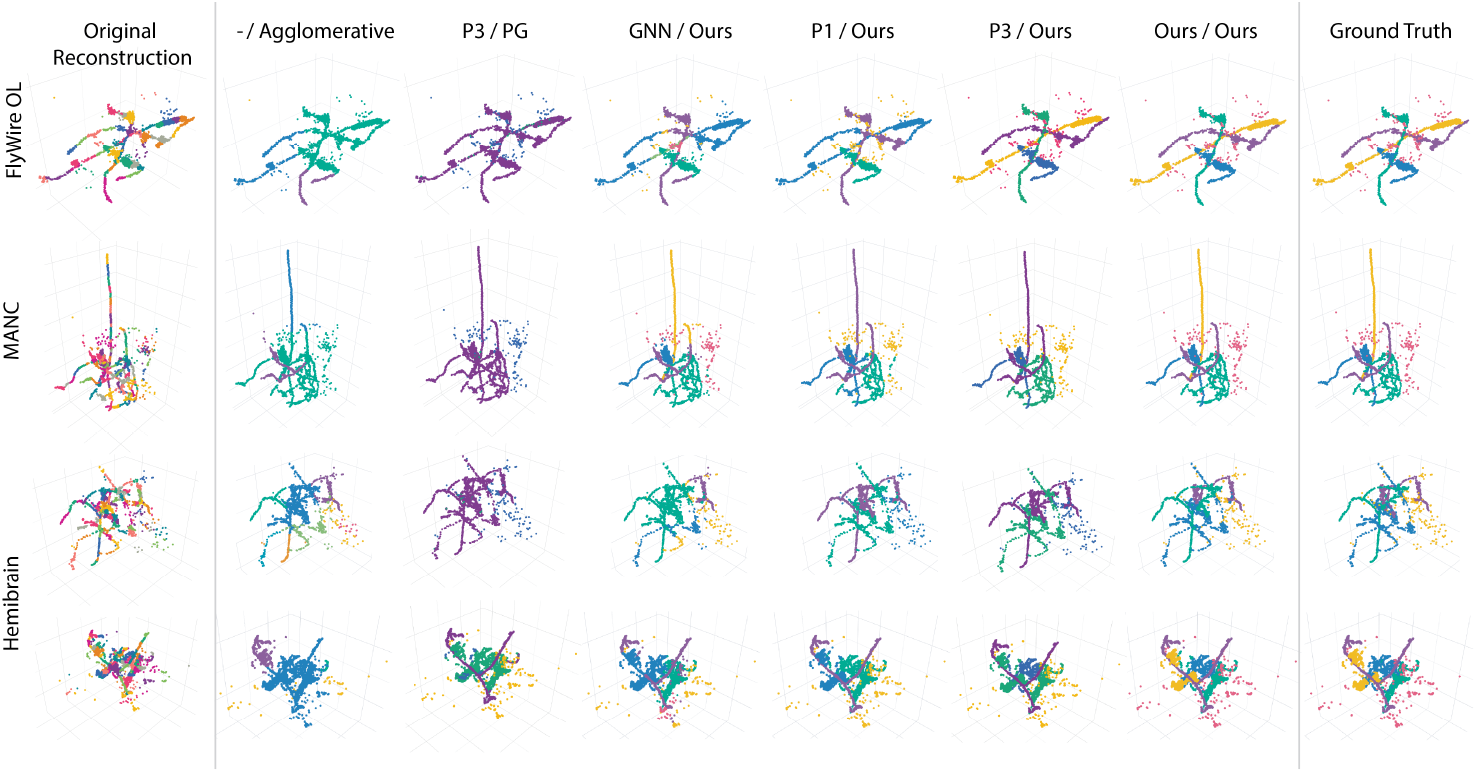
Automated Error Correction Examples. Our approach can reliably correct merge and split errors in neuron point clouds. Qualitative results across three datasets demonstrate our model’s performance. We compare our approach against unsupervised clustering methods such as agglomerative clustering, a graph neural network-based approach (Point-GNN), and the commonly used Point-Transformer V1 and V3 (P1 and P3) architecture. Baselines are named as (Encoder/Cluster Decoding). The color represents segmentation labels *S*.

We use feature agglomerative hierarchical clustering [49] to iteratively merge points into clusters according to 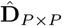 *P* _*×*_ *P*, an average linkage criterion and a termination threshold *t* (Fig. 2 d). Upon convergence of the agglomerative clustering algorithm, all points in a cluster get the same segmentation label *s* ∈ *S* assigned. This approach allows accurate reconstruction of segmentation labels, even if affinity predictions are imperfect. The background class (*s* = 0) is defined by the union of all clusters with less than 30 points. We find that 30 is a good threshold to summarize background fragments into the background class.

### 3.4 Contrastive Feature Embeddings

We use neuron type classification to demonstrate that our model learns to represent global neuron shapes, as the shape is among the most important contributing factors for neuron type. Given a pretrained affinity prediction model *M* (Sec. 3.1) and a single neuron point cloud *P*_*s*_, we retrieve the respective feature set 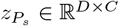 from *M* (Sec. 3.2). Next, we train a Deep Set [53] *g* : R *D×C* → R*E* that maps 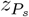 into a lower dimensional contrastive embedding space with dimensionality *E*. The Deep Set *g* consists of a feature embedding MLP *ζ*, a permutation invariant summation operation Σ and an output MLP *ρ* that projects into the contrastive embedding space:

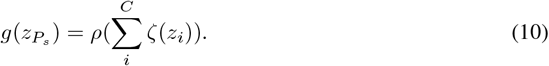

Additionally, we normalize contrastive embeddings, such that 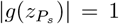. While keeping all parameters of *M* frozen when retrieving 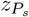, we train *g* with a contrastive loss function:

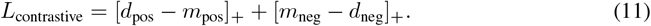

We use neuron type labels to compute Euclidean distances *d*_pos_ between embeddings derived from point clouds of neurons from the same type. *d*_neg_ are embedding distances between point clouds from different neuron types. *m*_pos_ and *m*_neg_ are the positive and negative margin values, respectively. Finally, an Euclidean KNN classifier computes neuron-type probabilities for single-neuron contrastive embeddings. Section 4.4 evaluates the performance of our contrastive embeddings.

**Table 1:** Automated Error Correction Evaluation. Our approach outperforms baselines across all three datasets in terms of the VOI, including the split VOI_*s*_ and merge VOI_*m*_ variants, and the adjusted rand error (ARE). The best scores are **bolded**, and the second-best scores are underlined. Baselines are named as (Encoder/Cluster Decoding). See Section 4.2 for details about baselines.

| Data | Metric | Agg. | P3/PG | GNN/Ours | P1/Ours | P3/Ours | Ours/Ours |
| --- | --- | --- | --- | --- | --- | --- | --- |
| FlyWire [8] | VOI ↓ | 1.24 | 1.47 | 0.82 | <u>0.32</u> | 0.37 | **0.17** |
| | $\text{VOI}_s$ ↓ | 0.37 | 0.21 | 0.42 | **0.07** | <u>0.14</u> | **0.07** |
| | $\text{VOI}_m$ ↓ | 0.87 | 1.26 | 0.42 | 0.26 | <u>0.24</u> | **0.10** |
|  | ARE ↓ | 0.44 | 0.55 | 0.24 | <u>0.12</u> | <u>0.12</u> | **0.04** |
| MANC [44] | VOI ↓ | 0.97 | 1.44 | 0.51 | <u>0.27</u> | 0.29 | **0.20** |
| | $\text{VOI}_s$ ↓ | 0.32 | 0.12 | 0.20 | **0.07** | 0.09 | <u>0.08</u> |
| | $\text{VOI}_m$ ↓ | 0.65 | 1.32 | 0.31 | <u>0.2</u> | <u>0.20</u> | **0.13** |
|  | ARE ↓ | 0.3 | 0.52 | 0.15 | <u>0.09</u> | 0.10 | **0.05** |
| Hemibr. [38] | VOI ↓ | 1.06 | 1.42 | 0.77 | 0.39 | <u>0.35</u> | **0.23** |
| | $\text{VOI}_s$ ↓ | 0.48 | 0.31 | 0.39 | **0.09** | <u>0.11</u> | **0.09** |
| | $\text{VOI}_m$ ↓ | 0.58 | 1.11 | 0.38 | 0.30 | <u>0.23</u> | **0.14** |
|  | ARE ↓ | 0.35 | 0.49 | 0.23 | 0.14 | <u>0.11</u> | **0.05** |

## 4 Experiments

We evaluate our approach on three benchmark datasets, which are derived from proofread connectomes [10, 44, 38], allowing us to evaluate our model’s performance accurately. We report results for automatically correcting simulated reconstruction errors and demonstrate our model’s neuron shape reasoning capability using its feature representation for neuron type classification.

### 4.1 Data

#### Datasets

The *FlyWire* optic lobe (OL) [8] is part of the first-ever full adult Drosophila connectome and includes well-defined cell type families [34]. The *MANC* [44] connectome describes the neural wiring architecture of an adult male fruit fly’s ventral nerve cord, responsible for signaling motor output to the fly’s wings and legs and receiving sensory input. The *hemibrain* dataset [38] is a proofread connectome reconstruction of a fruit fly’s central brain, responsible for navigation and integrating visual information.

#### Data Construction

Currently, no standardized real-world datasets exist for benchmarking automated error correction methods. To address this gap, we created three synthetic benchmark datasets, derived programmatically from state-of-the-art connectomes. Our goal in constructing these datasets is to realistically replicate common errors and artifacts found in neuron segmentation tasks by combining several neurons and background segments into single point clouds, enabling robust training and evaluation. For each connectome, we randomly sample 15, 000 neurons to generate training data and 2, 000 neurons for testing. These neuronal point clouds are obtained by randomly sampling vertices from the publicly available neuron skeletons while discarding edge connectivity information.

Additionally, we incorporate small background fragments, which commonly emerge in CNN-based image segmentation pipelines as artifacts (e.g., spine heads or neuron arborizations), by adding randomly selected small neuron branches (length ≤ 8 µm). We randomly vary the number of merged neurons in the range of [1, …, 4] ∈ N. The supplemental material shows examples with more (≥ 4) input neurons (Fig. 6). In summary, the input point cloud to our model is constructed by combining a random set of neurons and background fragments into a single point cloud.

**Table 2:** Neuron Type Classification Results. Our contrastive embeddings significantly outperform DGCNN-based [48] embeddings for neuron type classification. DGCNN is a GNN-based approach. The best scores are **bolded**.

| Dataset | Metrics | DGCNN [48] | Ours |
| --- | --- | --- | --- |
| FlyWire [8] | mAP ↑ | 0.62 | **0.77** |
| | Top1 Error ↓ | 0.39 | **0.19** |
| | Top5 Error ↓ | 0.07 | **0.03** |
| Hemibrain [44] | mAP ↑ | 0.53 | **0.73** |
| | Top1 Error ↓ | 0.38 | **0.23** |
| | Top5 Error ↓ | 0.08 | **0.04** |

#### Data Augmentation

We apply a series of data augmentations to increase data diversity. Before combining neurons into a single point cloud, we first center each neuron around the origin, which helps the model to learn shapes rather than overfitting on global position. Next, we jitter each point by a random factor of [0, 1um] ∈ R and randomly rotate [0, 200°] and translate individual neurons. Finally, we merge randomly selected neurons and background fragments and apply a global scaling factor to ensure all coordinates are within [−1, 1].

### 4.2 Baseline Approaches

We compare against six baselines and report results for different encoder and cluster decoding methods. Baselines are named as (Encoder/Cluster Decoding).

#### Unsupervised Clustering

We compare our affinity based clustering (Sec. 3.3), to traditional unsupervised approaches such as agglomerative clustering [49] (Agg.). We first compute pairwise Euclidean distances between all point pairs *P × P* and predict neuron clusters with each respective algorithm. We obtain hyperparameters by performing a grid search on a small training dataset and picking the set of hyperparameters that yields the best VOI score.

#### GNNs

Graph-based approaches are a set of popular methods for point cloud processing and classification [26]. We compare our approach to Point-GNN [43] (GNN), a popular message-passing model based on graph convolution. This approach predicts a feature per point, which is decoded to object labels in the original paper. For affinity prediction, we add an MLP decoder to the model that decodes point feature pairs to affinity values (GNN/Ours). We reconstruct a radius graph from the point cloud using radius = 32 and a maximum of 64 neighbors per point.

#### Point Transformers

We compare to Point Transformer V1 (P1) [57] and Point Transformer V3 [50] (P3). These architectures perform step-wise local neighborhood aggregation and predict features per point. We compare against the original P3 instance segmentation approach that uses Point Group [19] for clustering (P3/PG). This led to poor performance due to PG’s reliance on object centroids for clustering, which are non-discriminative for tightly entangled neurons. We investigated if pairing P1 and P3 encoders with our affinity-based cluster decoding could rescue their poor performance (P1/Ours, P3/Ours).

#### Contrastive Embeddings

We compare our contrastive embeddings (Sec. 3.4) against a representative GNN called DGCNN [48]. This approach builds a graph from a point cloud by connecting nearby points with edges. DGCNN implements graph convolution and predicts a feature per single neuron point cloud *P*_*s*_. Following the training procedure of our DeepSet *g* (Sec. 3.4), we optimize DGCNN using the contrastive loss function *L*_contrastive_. In contrast to our method, which uses a pretrained encoder and only optimizes the deep set parameters, DGCNN is trained end-to-end (Sec. 3.4). We also considered benchmarking our approach against other conventional approaches like NBLAST [4]. We omitted comparisons with NBLAST since it takes the absolute location of neurons within a template brain into account, which is a different problem setting.

### 4.3 Implementation

#### Automated Error Correction

Each point cloud contains 4288 samples, which includes 1024 points per neuron and up to 6 background fragments, with each up to 32 points. We randomly sample {1, …, 4} neurons during affinity prediction training and testing. For cases where the number of neurons is *<* 4, zero padding is used to get an equal number of points per batch. We also report data ablation for OOD testing with *>* 4 neurons (Sec. 4.5). Generally, based on manual analysis of segmentation errors and the datasets used, we find ≤ 4 neurons to be a reasonable training distribution. The feature set *z*_*P*_ uses *C* = 512 channels, each with a dimensionality of *D* = 1024. The encoder *φ* includes a series of 24 self-attention modules. The decoder’s MLP has two layers, each 2*D* = 2048 units. The first layer uses ReLU activations, and the output layer uses a Sigmoid activation function. The convergence threshold for the agglomerative clustering algorithm is *t* = 0.8. All affinity prediction experiments are trained for 946 epochs and a batch size of 230 on 8 NVIDIA H100 GPUs. For testing, we used a single H100 GPU. We use five warmup epochs to linearly increase the learning rate from 0 to *lr*_max_ = 1*e*^−4^ and use a cosine decay scheduler with *lr*_min_ = 1*e*^−6^.

**Figure 4:**
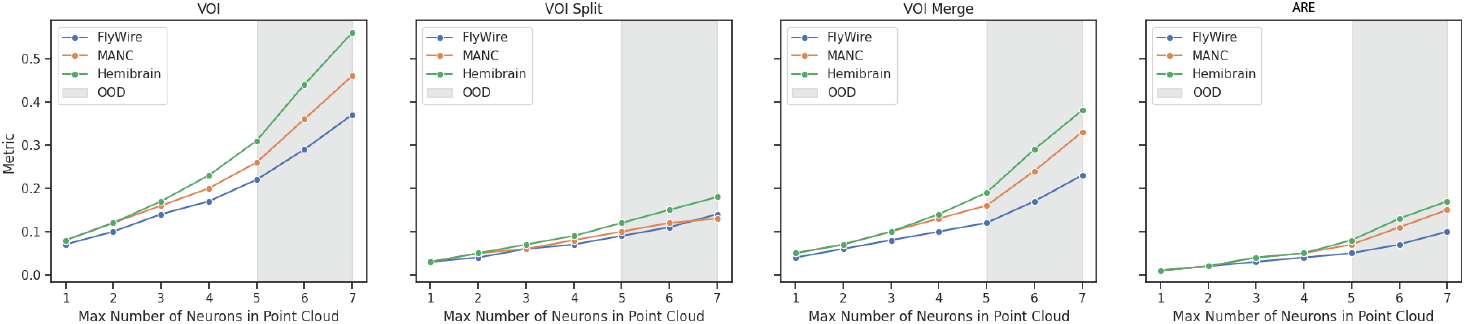
Input Data Size Ablation. We train our model with a maximum number of 4 neurons per point cloud. However, our model architecture is agnostic to input size. Hence, we report test set metrics for individual evaluations with a maximum of 1-7 neurons per point cloud. The grey area indicates out-of-distribution (OOD) performance. While we observe a slow linear decrease in-distribution (max 1-4 neurons), there is a bias toward under-segmentation OOD.

#### Neuron Type Classification

Here, we use single-neuron point clouds without background fragments, resulting in 1024 points. Using our pretrained encoder, the contrastive Deep Sets are trained for 100 epochs and a batch size of 650 on a single H100 GPU. The DGCNN baseline is trained end to end. The k-nearest neighbor classifier uses Euclidean distances and *k* = 15. For both experiments on FlyWire OL and the Hemibrain data (see Table. 4.4), we exclude neurons whose type has fewer than 40 instances in the respective dataset. Due to the imbalance of the neuron type distribution in both datasets, we rebalance the training dataset so that each type is equally represented. With these constraints, the Flywire training set has 35 unique types, each with 400 representative neurons. The hemibrain training dataset has 41 unique neuron types, each with 300 instances. The FlyWire test set includes around 33k neurons, while the hemibrain test set includes around 2600 neurons. The DeepSet MLP *ζ* and *ρ* consist of two linear layers with ReLU activations. *ζ* uses 1024 input units and 256 units in the hidden and output layers. *ρ* has 256 input and 32 output units with no activation function in the last layer. The contrastive embedding space has a dimensionality of *E* = 32.

All data, code, and trained models will be made accessible upon publication of this paper.

### 4.4 Main Results

#### Automated Error Correction

Table 1 shows quantitative comparisons of our approach to six baselines (Sec. 4.2). Our approach (Ours/Ours) consistently outperforms baselines. Broadly, learningbased approaches (GNNs, Transformers & Ours) outperform unsupervised clustering methods such as Agglomerative Clustering (Agg.). We also find that conventional instance segmentation techniques such as PTv3 with Point Group (P3/PG) [19] lead to poor performance, due to their reliance on object centroids for point clustering. However, combining the PTv3 encoder with our affinity based cluster decoding (P3/Ours) improves performance and achieves the second best scores in our experiments. Transformer architectures (P1/Ours, P3/Ours, Ours/Ours) are superior overall to GNNs (GNN/Ours). In contrast to Point Transformers, our method (Ours/Ours) produces significantly fewer neuron merge errors as shown in Table 1 (VOI_*m*_) and in Figure 3. This observation is consistent across all three benchmark datasets. We attribute these performance gains to our model’s ability to learn a feature representation without explicit spatial references, allowing flexible fitting to thin and sparse spatial objects, such as neurons. Our method depends on a threshold *t*, which determines the convergence of the agglomerative affinity clustering (Sec. 3.3). We created a small training set for each method and identified *t* with the highest VOI through a grid search.

#### Neuron Type Classification

Table 2 shows neuron type classification metrics for both our contrastive embeddings (Sec. 3.4) and the embeddings from our GNN-based baseline DGCNN [48] (Sec. 4.2). Our approach outperforms the baselines for both datasets on all reported metrics. Specifically, the Top1 error metric improved, with a −0.2 for FlyWire and a −0.15 for the Hemibrain over the baseline.

**Table 3:**
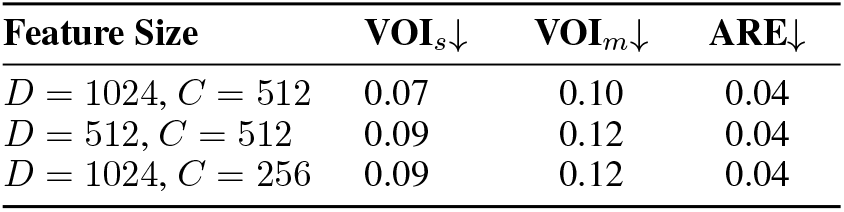
Feature Size Ablation. We ablate the size of the feature dimensionality *D* and the number of elements *C* in the feature set. Reducing both *D* or *C* equally degrades performance metrics.

| Feature Size | VOI<sub>s</sub>↓ | VOI<sub>m</sub>↓ | ARE↓ |
| --- | --- | --- | --- |
| $D = 1024, C = 512$ | 0.07 | 0.10 | 0.04 |
| $D = 512, C = 512$ | 0.09 | 0.12 | 0.04 |
| $D = 1024, C = 256$ | 0.09 | 0.12 | 0.04 |

### 4.5 Ablations

We report two ablation studies for the automated error correction task to answer the following questions:

- Question 1: Does our approach generalize to out-of-training distributions regarding the maximum number of neurons merged into a single point cloud?
- Question 2: Does a lower feature dimensionality *D & C* reduce performance numbers?

#### OOD Data Ablation (Q1)

We show our model performance on various numbers of neurons in a single point cloud for the automated error correction task (Fig 4). Our model was trained for only a maximum number of four neurons per point cloud. Thus, the grey area indicates OOD performance. As expected, we observe a linear performance decrease in-distribution (max. 1-4 neurons per point cloud). For OOD evaluations (5 - 7 neurons per point cloud), we identify a bias towards undersegmenting point clouds. In comparison, the VOI_*s*_ increases linearly for OOD input data, and VOI_*m*_ starts increasing more rapidly. This effect could be mitigated by decreasing the convergence threshold *t* of the affinity-based agglomerative clustering (Sec. 3.3).

#### Feature Set Size (Q2)

Table 3 shows that reducing either feature dimensionality *D* or the number of features *C* equally decreases VOI_*s*_ and VOI_*m*_. The ARE error metric does not decrease.

## 5 Discussion

The lack of neural network architectures capable of reasoning with the global shapes of neurons has been a major bottleneck for the complete automation of connectome generation. Solving this problem will enable connectome mapping of entire fly brains in mere weeks instead of years, making connectome mapping of entire mouse brains feasible. Our results show significant performance gains for both tasks over various baseline methods and datasets.

### Limitations

Here, we employed simulated neuron reconstruction errors to evaluate our automated error correction framework. This choice was motivated by two main factors. First, it enabled targeted testing in challenging scenarios, including varying numbers of input neurons (Fig. 4 & 6b) and cases where global shape alone may be ambiguous (Fig. 6a). Second, there is currently no standardized benchmark dataset available, due to the absence of matched pre- and post-proofreading versions of large-scale segmentation volumes.

### Broader Impact

Our framework, which leverages global neuron shapes, promises better generalization of pretrained models across connectomes from various imaging modalities, significantly speeding up the reconstruction process. This will accelerate scientific discovery in related fields and potentially enhance diagnostics and treatments for brain-related diseases. Potential negative impacts, such as misuse in cognitive profiling, remain a concern, albeit not a realistic threat in the near future.

### Future Work

Our work can be extended in several ways. Point clouds could be augmented with additional feature vectors based on local image information, for instance, based on SegCLR [7] or synapse type [11]. Neuron embeddings generated by our general-purpose encoder based on a small number of annotations in one brain region or animal species could then be used for the unsupervised discovery of new cell types in other brain regions and species. Our method for proofreading can be extended to tracing neurons in light microscopic images by proofreading point clouds segmented with simple binary thresholding. Finally, decoders could be trained to generate skeleton graph representations and computationally efficient renderings (Gaussian splatting [22]) of neuronal shapes from sparsely sampled point clouds.

## A Evaluation Metrics

### Automated Error Correction

We solve an affinity-guided clustering problem, where points of the same neuron form a cluster. Thus, we use standard clustering metrics such as the *variation of information* (VOI) and the *adjusted rand error* (ARE). Additionally, the VOI is decomposed into a split (VOI_*s*_) and merge (VOI_*m*_) component, which measures the degree of over- and under-segmentation, respectively.

**Figure 5:**
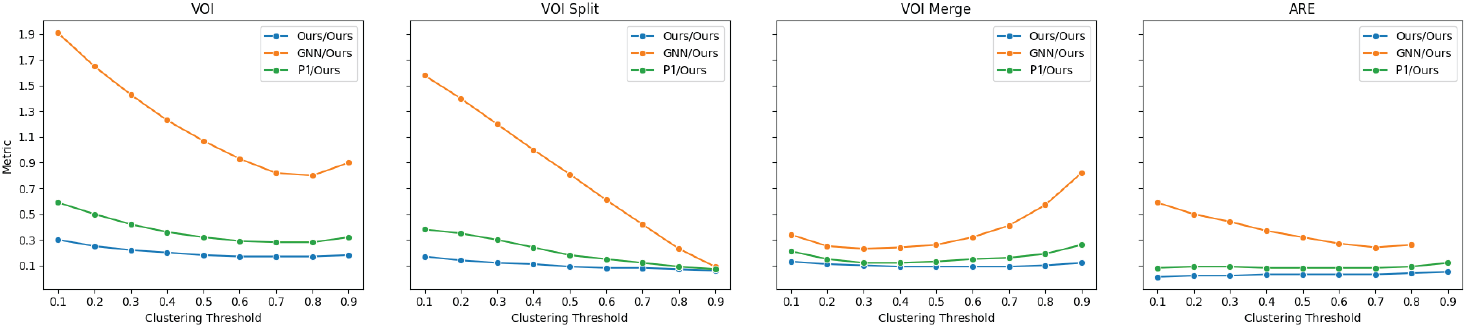
Clustering Threshold Ablation. These charts plot the agglomeration clustering threshold *t* (Sec. 3.3) relative to automated proofreading performance metrics for the FlyWire dataset. Here, we focus on learning-based approaches like GNN/Ours [43], P1/Ours [57], and our proposed approach. While performance fluctuations occur as the clustering threshold changes, our method consistently performs better than the reported baselines. All plots here are based on the FlyWire dataset.

### Neuron Type Classification

For this multi-class classification problem, we report mean average precision (*mAP*), the Top1 Error, and the Top5 Error. The Top1 error indicates the likelihood that the correct type is *not* the top prediction. The Top5 Error measures the likelihood that the correct neuron type is *not* in the top 5 predictions.

## B Additional Ablations

Figure 5 illustrates the sensitivity of our method and the respective baselines to the convergence threshold of the affinity-guided agglomeration clustering (Sec. 3.3). Our method outperforms baselines across all possible thresholds in terms of the variation of information (VOI) metric, its split (VOI_*s*_), and merges (VOI_*m*_) decompositions, and the adjusted rand error (ARAND). For all metrics, lower values indicate better performance.

## C Additional Qualitative Results

### Automated Error Correction

We provide additional qualitative results for the automatic proofreading use case. Figure 7 and Figure. 8 show examples for single-, double-, three- and four-neuron point clouds. Irrespective of the number of neurons, our approach outperforms the respective baselines. Notably, our approach reliably avoids oversegmenting single neuron point clouds (Fig 7).

### Results with Parallel Fibers

Figure 6a shows additional qualitative results for particularly challenging automated proofreading cases involving parallel neuronal fibers. We show a success and failure case (top/bottom row).

### More Input Neurons

The number of merged neurons depends on the agglomeration strategy applied to the CNN-based volumetric segmentation, and is thus a hyperparameter in our framework. Figure 6b shows additional qualitative results for 7 and 8 input neurons.

### Neuron Type Classification

Figure 10 shows the detailed confusion matrices for quantitative results reported in Table 11. The numbers in each tile represent the percentage (%) of neuron instances of a specific ground truth type, which were classified respectively. Additionally, Figure 9 shows three examples of different neuron morphologies. For each neuron, we also show the Top3 predicted neuron types based on our classifier (Sec. 3.4). The ground truth type is highlighted in green. For all three examples, the correct type is within the top two predicted neuron types.

**Table 4:** **Field of View (FOV) Comparison** for learning based automated error correction methods. The FOV for other approaches was estimated, given the parameter configuration in the respective papers. ^*^ Tested with up to 3*k ×* 8*k ×* 10*k* voxels.

| Method | FOV (voxels) | Architecture | Data Representation |
| --- | --- | --- | --- |
| Dmitriev [6] | $129 \times 129 \times 31$ | CNN | Skeleton + 3D Volume |
| Zung [58] | $136 \times 136 \times 136$ | CNN | 3D Volume |
| Haehn [16] | $46 \times 46 \times 1$ | CNN | 2D Image |
| Schmidt [16] | $33 \times 33 \times 11$ | CNN | 3D Volume |
| Ours | not limited * | Transformer | 3D Point Cloud |

## D Details for Baseline Implementations

We use the default hyperparameter configuration for the Point-GNN [43] (GNN), PT [57] (P1), PTv3 [50] (P3), PG [19] and DGCNN [48] baselines, as reported in the respective papers. The PTv3 encoder additionally uses a grid size parameter of 0.001 to achieve the best performance with our data. All affinity decoding models were trained using the binary cross-entropy loss. The PT baseline uses a convergence threshold for the agglomeration clustering of *t* = 0.9, PTv3 uses *t* = 0.8, and Point-GNN uses *t* = 0.7. We determined those parameters using grid search on a small training set. The unsupervised clustering baselines (Agg.) use a threshold of *t* = 0.1 and *t* = 0.3, respectively. All learning-based baselines were trained until the respective loss converged.

## E Scalability

Our model has a common point cloud encoder for both tasks and a task-dependent decoder. The encoder scales linearly with point cloud size through a constant-sized feature embedding (Fig. 2b). The neuron typing decoder scales constantly with point cloud size. The affinity decoder naively scales quadratically with point cloud size. However, strategies that decode affinities between supervoxels only require computation scaling quadratically with supervoxels, not points. For instance, with 4, 288 input points, decoding affinities between 2, 000 supervoxels only takes 70 ms on one H100 GPU. For extremely large inputs, such as 30, 000 points grouped into 7, 500 supervoxels, inference takes around 750 ms using one H100 GPU. Additional strategies for improved scalability include decoding affinities only locally in regions of interest during inference or decoding affinities with respect to a set of points of interest.

## F Alternative Neuron Error Correction Methods

Here, we briefly review existing approaches to neuron reconstruction error correction and highlight how our method differs from these alternatives.

### Heuristic Error Correction

Joyce et al. [20] employ handcrafted heuristics to identify potential 3D locations of reconstruction errors, focusing on the abrupt termination of neuronal branches, which often signals a split error. While effective for this specific error type, their method does not generalize to other error classes, most notably, merge errors. Celii et al. [2] decompose neuronal meshes to graphs and apply a set of heuristic rules to detect and correct merge errors. Similarly, Matejek et al. [33] incorporate biologically motivated geometric constraints to reconnect erroneously split segments. These methods depend on manually crafted priors about neuronal morphology, which may not adequately capture the full variability of reconstruction errors across datasets and brain regions.

### Learning-based Local Error Correction

Dmitriev et al. [6] use deep convolutional neural networks to detect split and merge errors in local skeleton graph neighborhoods. Haehn et al. [16] train a ConvNet to identify segmentation errors directly in 2D segmentation and image slices. Zung et al. [58] treat error correction as a pruning task over local volumetric segmentation masks. Schmidt et al. [39] approach the problem as a 3D flight tracing, where an agent guided by a CNN traverses neuronal branches to detect anomalies. All of these methods operate within restricted local fields of view, which limits their ability to resolve ambiguities that require broader spatial context.

**Figure 6:**
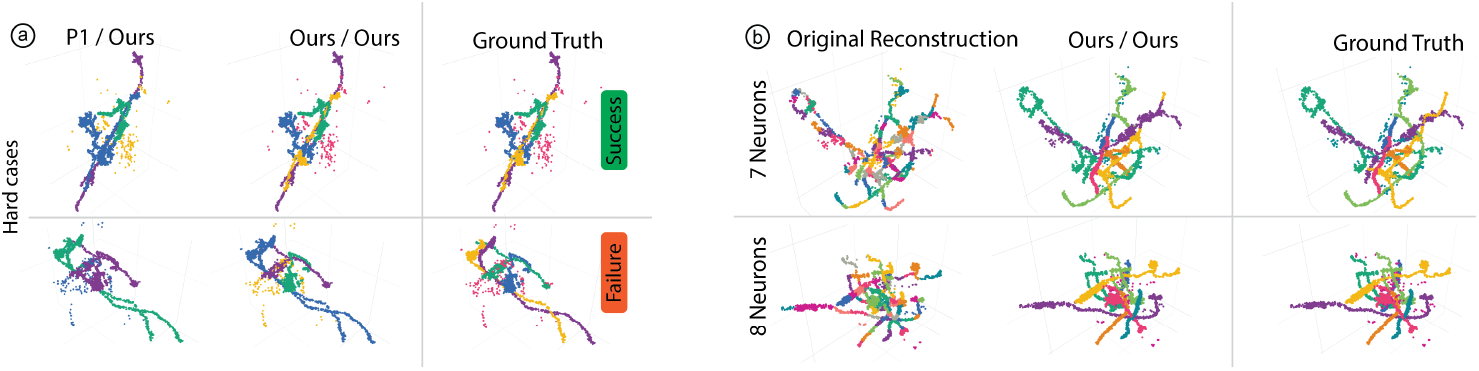
More qualitative results. (a) Results involving parallel fibers. The top row shows a success case. The bottom row shows a failure case. (b) Examples with 7 / 8 input neurons.

**Figure 7:**
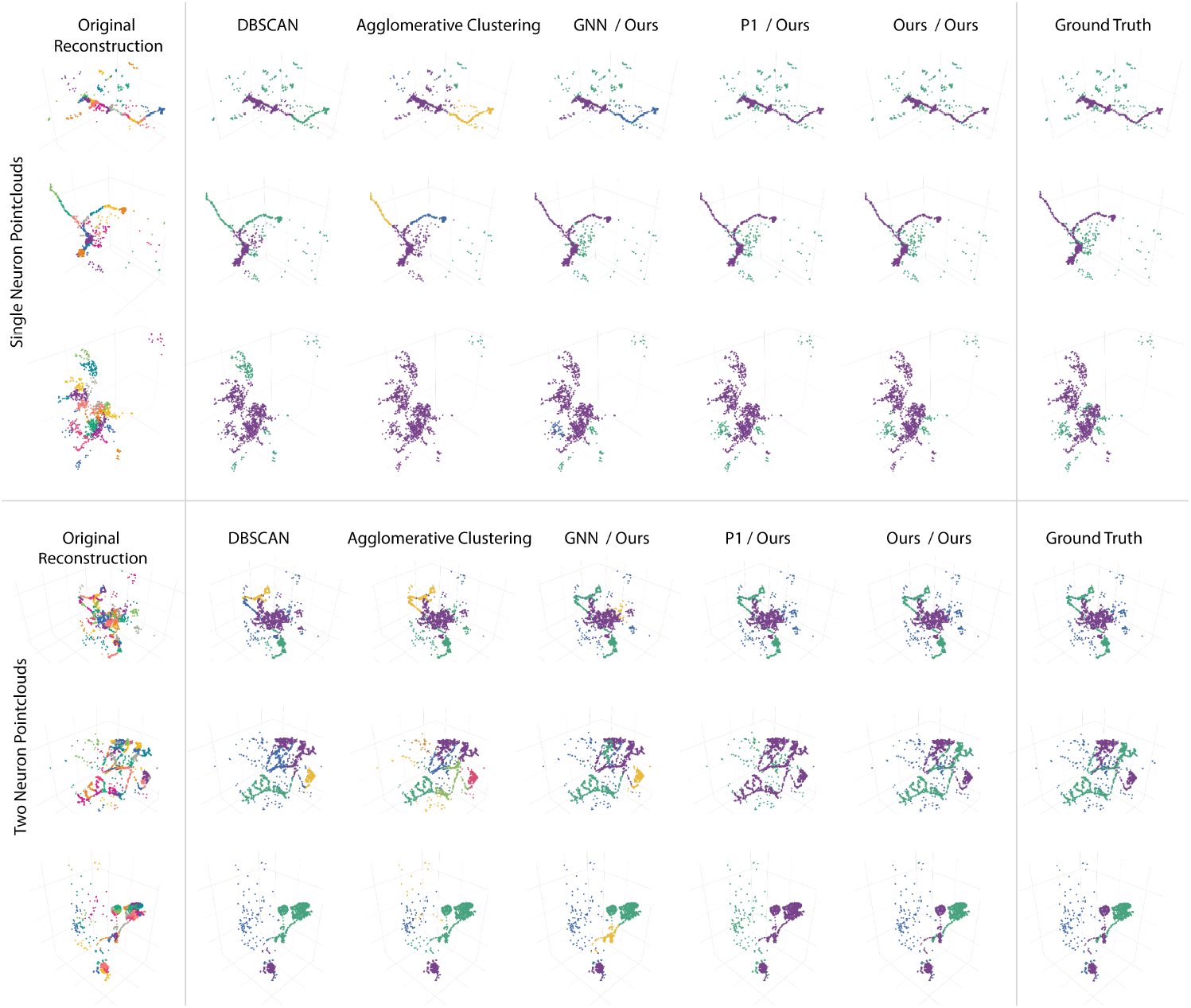
Additional Qualitative Results for Single & Two Neuron Pointclouds. We show additional examples for single neuron point clouds (top half) and two neuron point clouds (bottom half). Our approach reliably detects single neurons and does not oversegment them. Qualitative examples also show that our approach is more accurate for two-neuron point clouds than the shown baselines.

**Figure 8:**
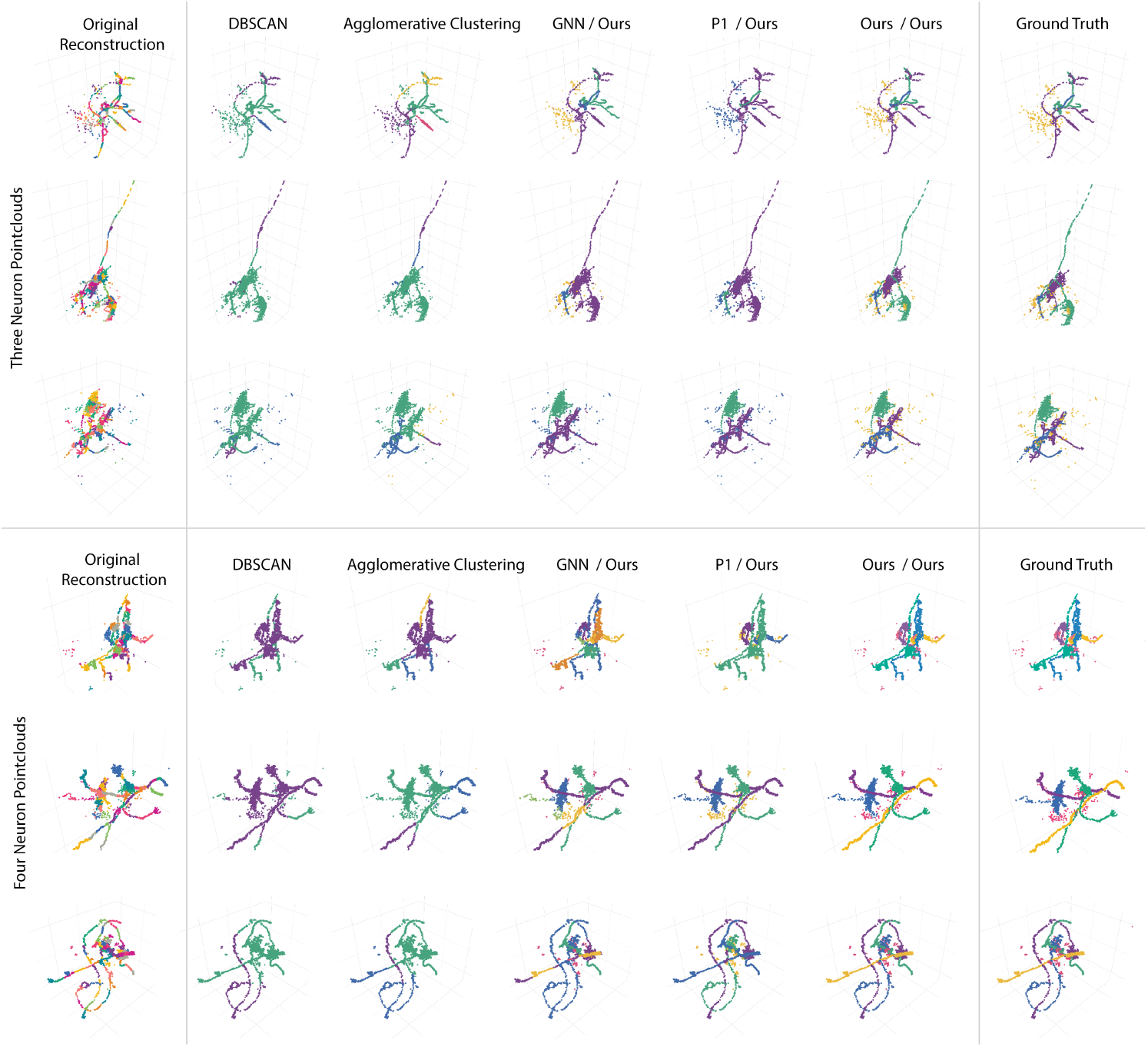
Additional Qualitative Results for Three & Four Neuron Point Clouds. The top half of the figure shows three neuron point clouds, while the lower half shows point clouds with four neurons. Our approach reliably detects whole neuron morphologies and can segment the point cloud into individual neurons. Notably, most baselines struggle with undersegmentation.

**Figure 9:**
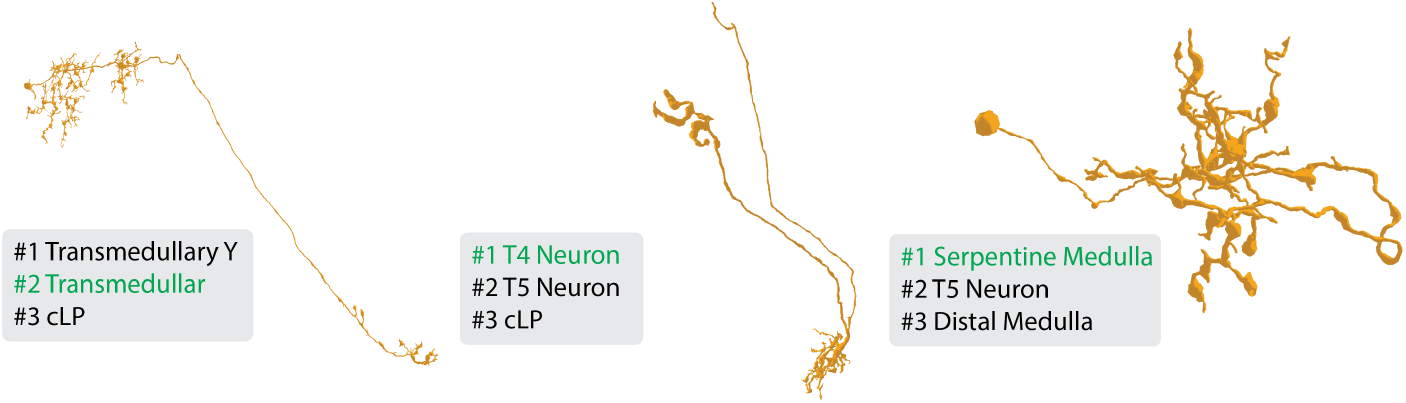
Neuron Type Examples & Classifier Predictions. We show three exemplary neuron morphologies and their Top3 predicted types using our KNN classifier. The ground truth type is shown in green. The correct type is within the top 2 predictions in all three examples. Table. 2 and Section 3.4 show more details and results.

**Figure 10:**
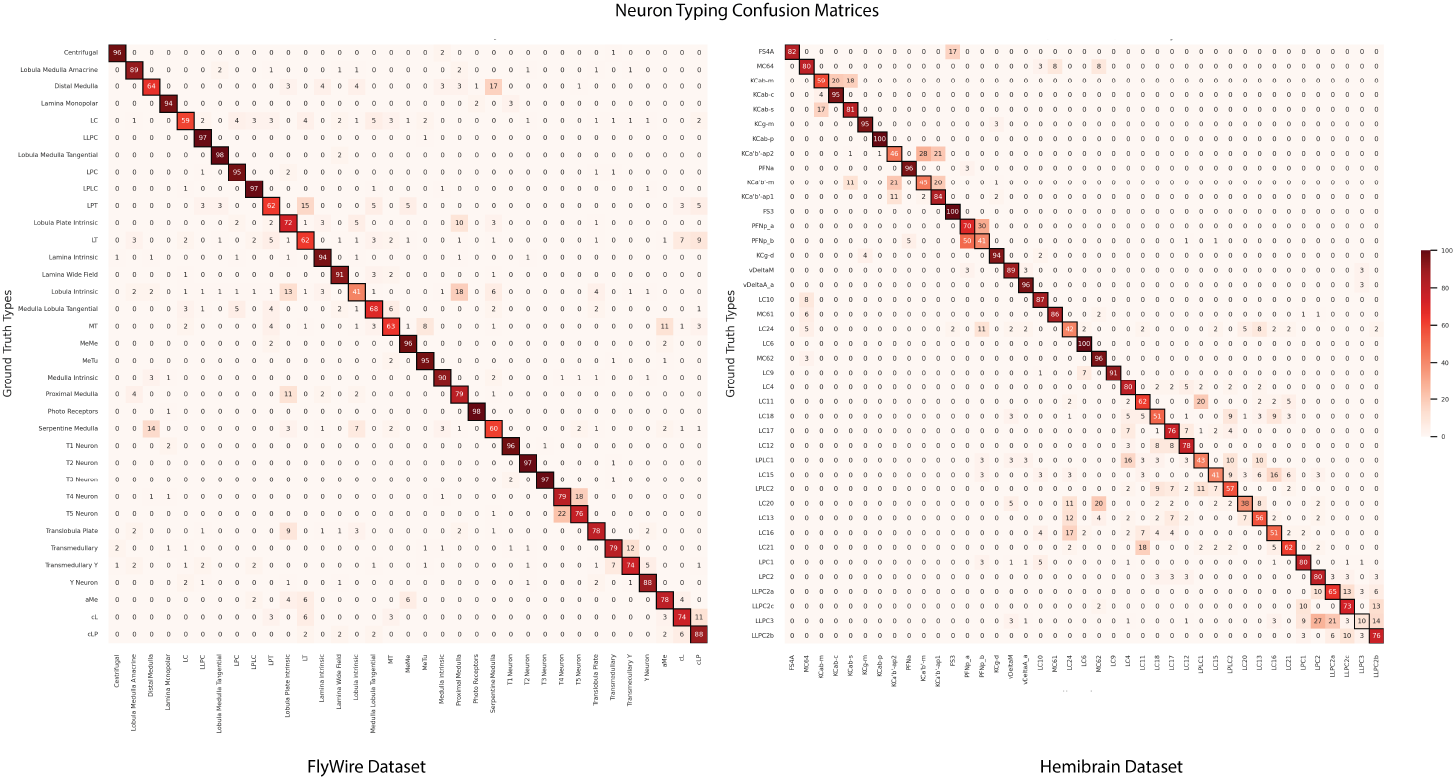
Neuron Typing Confusion Matrices. Here, we show neuron type classification confusion matrices for the FlyWire OL and the Hemibrain dataset. Both matrices are normalized, and percentage values (%) are shown in each matrix tile. The x-axis represents predicted types, while the y-axis represents ground truth types. More details are shown in Section 3.4.

**Figure 11:**
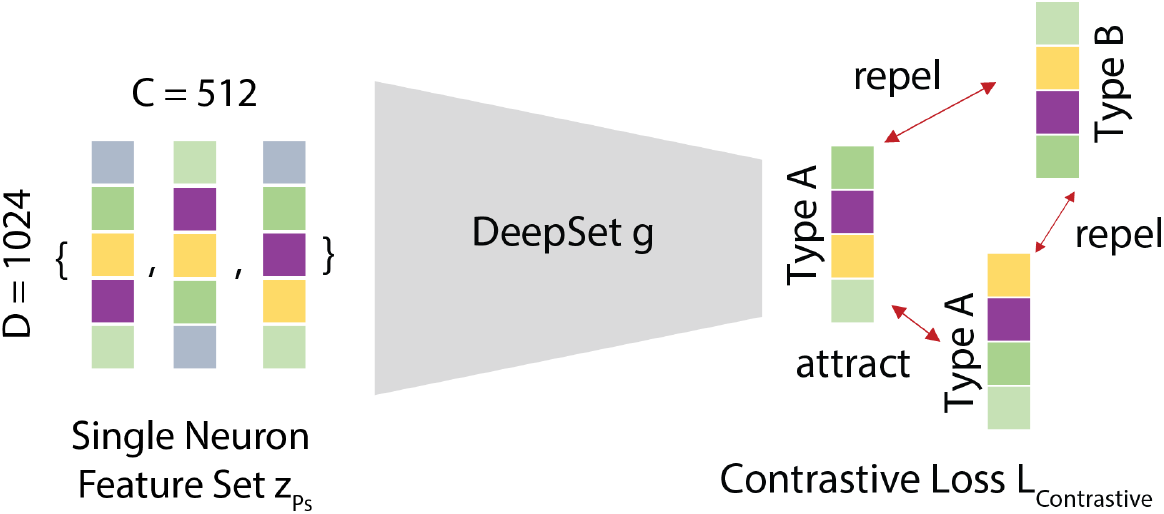
Neuron Type Contrastive Embeddings. We project a single neuron feature set *z*_*P*_*s* into a lower dimensional embedding space using a DeepSet *g*, trained using a contrastive loss function *L*_contrastive_. In the embedding space, neuron types are predicted with a k-nearest-neighbor classifier using Euclidean distances.

## Notes

### Competing Interest Statement

The authors have declared no competing interest.

### Summary of Updates

* updated figures, experiments & text.

## References

[1] Larry F Abbott, Davi D Bock, Edward M Callaway, Winfried Denk, Catherine Dulac, Adrienne L Fairhall, Ila Fiete, Kristen M Harris, Moritz Helmstaedter, Viren Jain, and others. The mind of a mouse. Cell, 182(6):1372–1376, 2020.

[2] Brendan Celii, Stelios Papadopoulos, Zhuokun Ding, Paul G Fahey, Eric Wang, Christos Papadopoulos, Alexander B Kunin, Saumil Patel, J Alexander Bae, Agnes L Bodor, et al. Neurd offers automated proofreading and feature extraction for connectomics. Nature, 640(8058):487–496, 2025.

[3] Hanbo Chen, Jiawei Yang, Daniel Maxim Iascone, Lijuan Liu, Lei He, Hanchuan Peng, and Jianhua Yao. TreeMoCo: contrastive neuron morphology representation learning. In Proceedings of the 36th International Conference on Neural Information Processing Systems, NIPS ‘22, Red Hook, NY, USA, 2024.

[4] Marta Costa, James D Manton, Aaron D Ostrovsky, Steffen Prohaska, and Gregory SXE Jefferis. NBLAST: rapid, sensitive comparison of neuronal structure and construction of neuron family databases. Neuron, 91(2):293–311, 2016.

[5] Konstantin Dmitriev12, Toufiq Parag, Brian Matejek, Arie E Kaufman12, and Hanspeter Pfister. Efficient correction for em connectomics with skeletal representation. British Machine Vision Conferemce (BMVC), 2018.

[6] Konstantin Dmitriev12, Toufiq Parag, Brian Matejek, Arie E Kaufman12, and Hanspeter Pfister. Efficient correction for em connectomics with skeletal representation. British Machine Vision Conferemce (BMVC), 2018.

[7] Sven Dorkenwald, Peter H Li, Michał Januszewski Daniel R Berger, Jeremy Maitin-Shepard, Agnes L Bodor, Forrest Collman, Casey M Schneider-Mizell, Nuno Maçarico da Costa, Jeff W Lichtman, and others. Multi-layered maps of neuropil with segmentation-guided contrastive learning. Nature Methods, 20(12):2011–2020, 2023.

[8] Sven Dorkenwald, Arie Matsliah, Amy R Sterling, Philipp Schlegel, Szi-Chieh Yu, Claire E McKellar, Albert Lin, Marta Costa, Katharina Eichler, Yijie Yin, and others. Neuronal wiring diagram of an adult brain. Nature, 634(8032):124–138, 2024.

[9] Sven Dorkenwald, Claire E McKellar, Thomas Macrina, Nico Kemnitz, Kisuk Lee, Ran Lu, Jingpeng Wu, Sergiy Popovych, Eric Mitchell, Barak Nehoran, and others. FlyWire: online community for whole-brain connectomics. Nature methods, 19(1):119–128, 2022.

[10] Sven Dorkenwald, Casey M Schneider-Mizell, Derrick Brittain, Akhilesh Halageri, Chris Jordan, Nico Kemnitz, Manual A Castro, William Silversmith, Jeremy Maitin-Shephard, Jakob Troidl, and others. CAVE: Connectome annotation versioning engine. bioRxiv, 2023.

[11] Nils Eckstein, Alexander Shakeel Bates, Andrew Champion, Michelle Du, and et al. Neurotransmitter classification from electron microscopy images at synaptic sites in Drosophila melanogaster. Cell, 187(10):2574–2594.e23, May 2024.

[12] Yimin Fan, Yaxuan Li, Yunhua Zhong, Liang Hong, Lei Li, and Yu Li. Learning meaningful representation of single-neuron morphology via large-scale pre-training. Bioinformatics, 40(Supplement_2):ii128–ii136, September 2024.

[13] Jan Funke, Fabian Tschopp, William Grisaitis, Arlo Sheridan, Chandan Singh, Stephan Saalfeld, and Srinivas C Turaga. Large scale image segmentation with structured loss based deep learning for connectome reconstruction. IEEE transactions on pattern analysis and machine intelligence, 41(7):1669–1680, 2018.

[14] Benjamin Gallusser and Martin Weigert. Trackastra: Transformer-based cell tracking for live-cell microscopy. In European Conference on Computer Vision, pages 467–484. Springer, 2024.

[15] Felix Gonda, Xueying Wang, Johanna Beyer, Markus Hadwiger, Jeff W Lichtman, and Hanspeter Pfister. VICE: Visual identification and correction of neural circuit errors. In Computer graphics forum, volume 40, pages 447–458, 2021. Issue: 3.

[16] Daniel Haehn, Verena Kaynig, James Tompkin, Jeff W Lichtman, and Hanspeter Pfister. Guided proofreading of automatic segmentations for connectomics. In Proceedings of the IEEE Conference on Computer Vision and Pattern Recognition, pages 9319–9328, 2018.

[17] Michał Januszewski Jörgen Kornfeld, Peter H Li, Art Pope, Tim Blakely, Larry Lindsey, Jeremy Maitin-Shepard, Mike Tyka, Winfried Denk, and Viren Jain. High-precision automated reconstruction of neurons with flood-filling networks. Nature methods, 15(8):605–610, 2018.

[18] Jiaxiang Jiang, Michael Goebel, Cezar Borba, William Smith, and BS Manjunath. A robust approach to 3D neuron shape representation for quantification and classification. BMC bioinformatics, 24(1):366, 2023.

[19] Li Jiang, Hengshuang Zhao, Shaoshuai Shi, Shu Liu, Chi-Wing Fu, and Jiaya Jia. Pointgroup: Dual-set point grouping for 3d instance segmentation. In Proceedings of the IEEE/CVF conference on computer vision and Pattern recognition, pages 4867–4876, 2020.

[20] Justin Joyce, Rupasri Chalavadi, Joey Chan, Sheel Tanna, Daniel Xenes, Nathanael Kuo, Victoria Rose, Jordan Matelsky, Lindsey Kitchell, Caitlyn Bishop, and others. A Novel Semi-automated Proofreading and Mesh Error Detection Pipeline for Neuron Extension. bioRxiv, 2023.

[21] Lida Kanari, Paweł Dłotko, Martina Scolamiero, Ran Levi, Julian Shillcock, Kathryn Hess, and Henry Markram. A topological representation of branching neuronal morphologies. Neuroinformatics, 16:3–13, 2018. Publisher: Springer.

[22] Bernhard Kerbl, Georgios Kopanas, Thomas Leimkühler, and George Drettakis. 3D Gaussian Splatting for Real-Time Radiance Field Rendering. 2023.

[23] Reem Khalil, Sadok Kallel, Ahmad Farhat, and Pawel Dlotko. Topological sholl descriptors for neuronal clustering and classification. PLOS Computational Biology, 18(6):e1010229, 2022.

[24] Longxin Le and Yimin Wang. 3D-aware neural network for analyzing neuron morphology. In 2024 5th International Conference on Computer Engineering and Application (ICCEA), pages 101–104, Hangzhou, China, April 2024. IEEE.

[25] Kisuk Lee, Jonathan Zung, Peter Li, Viren Jain, and H. Sebastian Seung. Superhuman Accuracy on the SNEMI3D Connectomics Challenge, 2017. Version Number: 1.

[26] Dilong Li, Chenghui Lu, Ziyi Chen, Jianlong Guan, Jing Zhao, and Jixiang Du. Graph Neural Networks in Point Clouds: A Survey. Remote Sensing, 16(14):2518, 2024.

[27] Hanyu Li, Michał Januszewski Viren Jain, and Peter H Li. Neuronal subcompartment classification and merge error correction. In Medical Image Computing and Computer Assisted Intervention–MICCAI 2020: 23rd International Conference, Lima, Peru, October 4–8, 2020, Proceedings, Part V 23, pages 88–98. Springer, 2020.

[28] Minghui Liao, Guojia Wan, and Bo Du. Joint Learning Neuronal Skeleton and Brain Circuit Topology with Permutation Invariant Encoders for Neuron Classification. In AAAI, volume 38, pages 197–205, 2024. Issue: 1.

[29] Zudi Lin, Donglai Wei, Aarush Gupta, Xingyu Liu, Deqing Sun, and Hanspeter Pfister. Structure-Preserving Instance Segmentation via Skeleton-Aware Distance Transform. In MICCAI, pages 529–539. Springer, 2023.

[30] Zudi Lin, Donglai Wei, Won-Dong Jang, Siyan Zhou, and others. Two stream active query suggestion for active learning in connectomics. In ECCV 2020, pages 103–120. Springer, 2020.

[31] Xiaoyu Liu, Miaomiao Cai, Yinda Chen, Yueyi Zhang, Te Shi, Ruobing Zhang, Xuejin Chen, and Zhiwei Xiong. Cross-dimension affinity distillation for 3d em neuron segmentation. In CVPR, pages 11104–11113, 2024.

[32] Yixiong Liu, Qihua Chen, and Xuejin Chen. Neuproofreader: An Interactive Proofreading System with Suggestive Prompts for Connectomics. In 2024 IEEE International Conference on Multimedia and Expo Workshops (ICMEW), pages 1–2. IEEE, 2024.

[33] Brian Matejek, Daniel Haehn, Haidong Zhu, Donglai Wei, Toufiq Parag, and Hanspeter Pfister. Biologically-constrained graphs for global connectomics reconstruction. In Proceedings of the IEEE/CVF conference on computer vision and pattern recognition, pages 2089–2098, 2019.

[34] Arie Matsliah, Szi-chieh Yu, Krzysztof Kruk, Doug Bland, Austin T Burke, Jay Gager, James Hebditch, Ben Silverman, Kyle Patrick Willie, Ryan Willie, and others. Neuronal parts list and wiring diagram for a visual system. Nature, 2024.

[35] Songyou Peng, Michael Niemeyer, Lars Mescheder, Marc Pollefeys, and Andreas Geiger. Convolutional occupancy networks. In ECCV, pages 523–540. Springer, 2020.

[36] Charles R Qi, Hao Su, Kaichun Mo, and Leonidas J Guibas. Pointnet: Deep learning on point sets for 3d classification and segmentation. In Proceedings of the IEEE CVPR, 2017.

[37] Charles Ruizhongtai Qi, Li Yi, Hao Su, and Leonidas J Guibas. Pointnet++: Deep hierarchical feature learning on point sets in a metric space. NeurIPS, 30, 2017.

[38] Louis K Scheffer, C Shan Xu, Michal Januszewski, Zhiyuan Lu, Shin-ya Takemura, Kenneth J Hayworth, Gary B Huang, and et al. A connectome and analysis of the adult Drosophila central brain. eLife, 9:e57443, September 2020.

[39] Martin Schmidt, Alessandro Motta, Meike Sievers, and Moritz Helmstaedter. RoboEM: automated 3D flight tracing for synaptic-resolution connectomics. Nature Methods, 21(5):908–913, 2024.

[40] Philipp J Schubert, Sven Dorkenwald, Michał Januszewski Viren Jain, and Joergen Kornfeld. Learning cellular morphology with neural networks. Nature communications, 10(1):2736, 2019.

[41] Alexander Shapson-Coe, Michał Januszewski Daniel R Berger, Art Pope, Yuelong Wu, Tim Blakely, Richard L Schalek, Peter H Li, Shuohong Wang, Jeremy Maitin-Shepard, and others. A petavoxel fragment of human cerebral cortex reconstructed at nanoscale resolution. Science, 384(6696):eadk4858, 2024.

[42] Arlo Sheridan, Tri M Nguyen, Diptodip Deb, Wei-Chung Allen Lee, Stephan Saalfeld, Srinivas C Turaga, Uri Manor, and Jan Funke. Local shape descriptors for neuron segmentation. Nature methods, 20(2):295– 303, 2023.

[43] Weijing Shi, Ragunathan, and Rajkumar. Point-GNN: Graph Neural Network for 3D Object Detection in a Point Cloud, 2020. Version Number: 1.

[44] Shin-ya Takemura, Kenneth J Hayworth, Gary B Huang, Michal Januszewski, Zhiyuan Lu, Elizabeth C Marin, Stephan Preibisch, C Shan Xu, John Bogovic, Andrew S Champion, and others. A connectome of the male drosophila ventral nerve cord. BioRxiv, pages 2023–06, 2023.

[45] Matthew Tancik, Pratul Srinivasan, Ben Mildenhall, Sara Fridovich-Keil, Nithin Raghavan, Utkarsh Singhal, Ravi Ramamoorthi, Jonathan Barron, and Ren Ng. Fourier features let networks learn high frequency functions in low dimensional domains. NeurIPS, 33:7537–7547, 2020.

[46] Hidetoshi Urakubo and Torsten Bullmann. UNI-EM: an environment for deep neural network-based automated segmentation of neuronal electron microscopic images. Scientific reports, 9(1), 2019.

[47] A Vaswani. Attention is all you need. Advances in Neural Information Processing Systems, 2017.

[48] Yue Wang, Yongbin Sun, Ziwei Liu, Sanjay E Sarma, Michael M Bronstein, and Justin M Solomon. Dynamic graph cnn for learning on point clouds. ACM Transactions on Graphics (tog), 38(5):1–12, 2019.

[49] Joe H Ward Jr. Hierarchical grouping to optimize an objective function. Journal of the American statistical association, 58(301):236–244, 1963. Publisher: Taylor & Francis.

[50] Xiaoyang Wu and et al. Point Transformer V3. In CVPR, 2024.

[51] Xiaoyang Wu, Yixing Lao, Li Jiang, Xihui Liu, and Hengshuang Zhao. Point transformer v2: Grouped vector attention and partition-based pooling. Advances in Neural Information Processing Systems, 35:33330– 33342, 2022.

[52] Feng Xiong, Peng Xie, Zuohan Zhao, Yiwei Li, Sujun Zhao, Linus Manubens-Gil, Lijuan Liu, and Hanchuan Peng. DSM: Deep sequential model for complete neuronal morphology representation and feature extraction. Patterns, 5(1):100896, January 2024.

[53] Manzil Zaheer, Satwik Kottur, Siamak Ravanbakhsh, Barnabas Poczos, Russ R Salakhutdinov, and Alexander J Smola. Deep sets. NeurIPS, 30, 2017.

[54] Biao Zhang, Matthias Nießner, and Peter Wonka. 3dilg: Irregular latent grids for 3d generative modeling. Advances in Neural Information Processing Systems, 2022.

[55] Biao Zhang, Jiapeng Tang, Matthias Niessner, and Peter Wonka. 3dshape2vecset: A 3d shape representation for neural fields and generative diffusion models. ACM Transactions on Graphics (TOG), 42(4):1–16, 2023.

[56] Cheng Zhang, Haocheng Wan, Xinyi Shen, and Zizhao Wu. Patchformer: An efficient point transformer with patch attention. In Proceedings of the IEEE/CVF CVPR, pages 11799–11808, 2022.

[57] Hengshuang Zhao, Li Jiang, Jiaya Jia, Philip Torr, and Vladlen Koltun. Point Transformer, 2020.

[58] Jonathan Zung, Ignacio Tartavull, Kisuk Lee, and H. Sebastian Seung. An Error Detection and Correction Framework for Connectomics, 2017. Version Number: 2.

